# Autocycler: long-read consensus assembly for bacterial genomes

**DOI:** 10.1101/2025.05.12.653612

**Authors:** Ryan R Wick, Benjamin P Howden, Timothy P Stinear

## Abstract

**Motivation:** Long-read sequencing enables complete bacterial genome assemblies, but individual assemblers are imperfect and often produce sequence-level and structural errors. Consensus assembly using Trycycler can improve accuracy, but its lack of automation limits scalability. There is a need for an automated method to generate high-quality consensus bacterial genome assemblies from long-read data.

**Results:** We present Autocycler, a command-line tool for generating accurate bacterial genome assemblies by combining multiple alternative long-read assemblies of the same genome. Without requiring user input, Autocycler builds a compacted De Bruijn graph from the input assemblies, clusters and filters contigs, trims overlaps and resolves consensus sequences by selecting the most common variant at each locus. It also supports manual curation when desired, allowing users to refine assemblies in challenging or important cases. In our evaluation using Oxford Nanopore Technologies reads from five bacterial isolates, Autocycler outperformed individual assemblers, automated pipelines and other consensus tools, producing assemblies with lower error rates and improved structural accuracy.

**Availability and implementation:** Autocycler is implemented in Rust, open-source and freely available at github.com/rrwick/Autocycler. It runs on Linux and macOS and is extensively documented.

## 1. Introduction

Complete genome assemblies are essential for resolving bacterial genome structure and fully characterising accessory elements such as plasmids and prophages. ^1,2^ Accurate assemblies reduce the risk of errors in downstream analyses such as comparative genomics, annotation and studies of genome dynamics.

Long-read sequencing platforms, such as those from Oxford Nanopore Technologies (ONT), have made complete assemblies of bacterial genomes widely achievable. Long reads can span repetitive elements, allowing assemblers to resolve structural complexity that short reads (e.g. from Illumina platforms) cannot. ^3^ For most bacterial genomes and high-quality read sets, long-read assemblers can assemble each replicon into a single contig. ^4^

In practice, however, long-read assemblers are imperfect, and different tools produce different assemblies from the same input read set. Common problems include: incomplete or overlapping circularisation, missing small plasmids, duplicated small plasmids and spurious extra contigs from repeats or contamination. ^5,6^ No single assembler is reliably the best across all datasets.

Consensus assembly offers a solution. By combining multiple alternative assemblies of the same genome (e.g. those produced by different assemblers or read subsets) consistent sequences can be distinguished from assembler-specific errors. ^7,8^ The software Trycycler put this idea into practice for bacterial genomes. ^9^ Compared to assemblies produced by a single tool, Trycycler assemblies usually contain fewer errors, more reliable circularisation and a more complete and less contaminated representation of the genome. ^9^

Trycycler, however, relies on human interventions and decision-making for several key steps. While this design offers flexibility and control, it limits scalability. As bacterial genomics increasingly involves large datasets of hundreds or thousands of genomes, there is a need for automated methods that can generate high-quality consensus assemblies without manual interventions.

Here we present Autocycler, a fully automated command-line tool to generate consensus long-read assemblies of bacterial genomes. Like Trycycler, it combines multiple input assemblies to produce a high-quality consensus. Unlike Trycycler, Autocycler is designed to run to completion without user input. It also supports manual intervention for cases where careful output curation is warranted. For most bacterial genomes and read sets with sufficient depth and read length, Autocycler can produce complete assemblies automatically. In more difficult cases, such as genomes with large repeats, genomic heterogeneity or unusual structures like linear replicons, users can step in to refine the output.

## 2. Implementation

Autocycler constructs a consensus bacterial genome assembly by combining multiple alternative assemblies of the same genome (Figure 1). It is designed to run fully automatically and produces intermediate files and metrics for every step, allowing the process to be inspected or curated if needed. In addition to the main consensus assembly workflow, Autocycler includes commands to assist with upstream and downstream tasks.

**Fig. 1.**
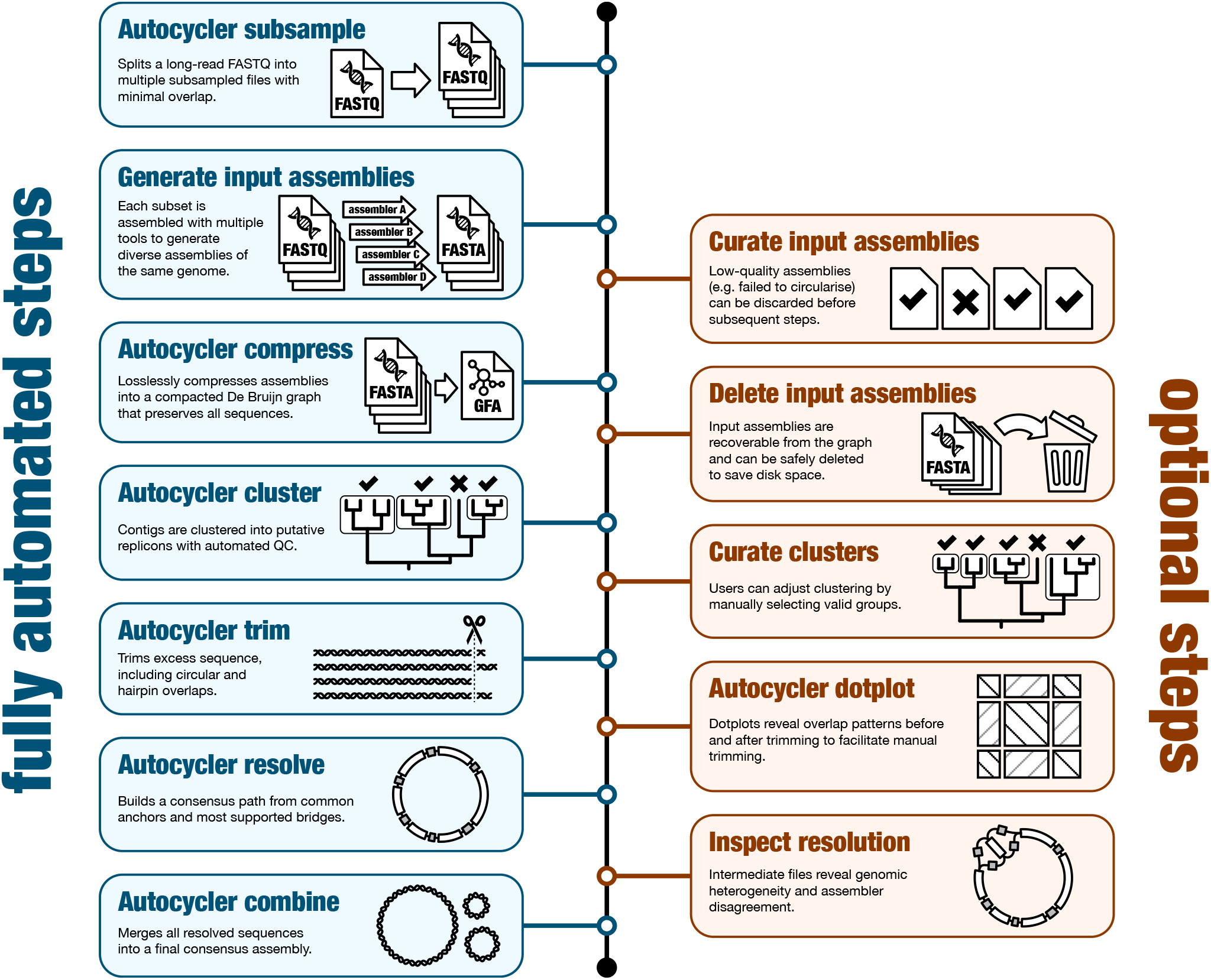
Overview of the Autocycler workflow. By following only the blue steps on the left, Autocycler can produce consensus genome assemblies with no human intervention. The optional orange steps on the right can be used when accuracy is critical or when Autocycler’s metrics (gathered using the `autocycler table` command) indicate potential issues.

The `autocycler subsample` command creates read subsets for generating input assemblies. By dividing a single long-read set into minimally overlapping subsets, users can generate assemblies that are more independent of one another. Additionally, the subsampling process reduces very high-depth read sets to medium-depth subsets, which often assemble more cleanly.

Input assemblies can be generated using any long-read assembler, and Autocycler provides helper scripts for several common ones. Ideally, each replicon in the genome (e.g. chromosome or plasmid) should be assembled as a single contig. While some fragmentation is tolerated, Autocycler relies on the assumption that most input assemblies are complete. If all input assemblies are fragmented, Autocycler will not be able to produce a complete consensus.

The `autocycler compress` command builds a compacted De Bruijn graph from the input assemblies. This graph requires much less disk space than the inputs, as shared regions are collapsed together. Importantly, it stores the full path of each input sequence, allowing the original assemblies to be recovered later using `autocycler decompress`. The graph representation also provides an efficient structure for manipulating and comparing sequences in downstream steps.

In the `autocycler cluster` step, input contigs are grouped based on similarity using UPGMA clustering. Autocycler then applies quality control filters to exclude low-confidence clusters (e.g. those with too few contigs or contained within other clusters), ideally leaving one high-quality cluster for each replicon in the genome. This step also outputs a Newick-format tree, which users can inspect to decide whether to override the automatic clustering.

For each cluster, `autocycler trim` processes the input contigs to remove unwanted sequence. It looks for both circular overlaps, where the start of a contig overlaps with its end, and hairpin overlaps, where the start or end of a contig extends past the hairpin to the opposite strand. It also handles cases where small plasmids are fully duplicated within a single contig. After trimming, sequences with lengths that deviate too far from the cluster median are discarded, leaving a set of consistent contigs for consensus generation.

The `autocycler resolve` command generates a consensus sequence for each cluster. It begins by identifying anchors: sequences that appear exactly once in each contig. These anchors serve as a scaffold for constructing bridges, which represent the most common paths between anchors in the input contigs. Autocycler first applies unambiguous bridges and then iteratively resolves ambiguous cases by selecting the most supported paths, ideally producing a single consensus sequence for the cluster. In cases of structural heterogeneity, such as phase-variable loci or assembly inconsistencies, Autocycler includes intermediate output for user inspection.

Once each cluster has been resolved, the `autocycler combine` command merges them into a final consensus assembly in both FASTA and GFA formats. Autocycler also produces detailed metrics at every step of the pipeline, saved in YAML format (both human- and machine-readable). The `autocycler table` command can be used to gather metrics from many assemblies, making it easy to track success and identify samples that require further attention.

Autocycler is implemented in Rust and is fast, deterministic and resource-efficient. Most of the computational time in an Autocycler workflow is spent generating input assemblies. Autocycler itself typically completes in minutes and requires only modest resources. It runs on both Linux and macOS, although Linux is preferred due to broader compatibility with long-read assemblers. Extensive documentation, including worked examples and guidance for manual curation, is available online at github.com/rrwick/Autocycler/wiki.

## 3. Evaluation

### 3.1 Methods

Long-read sequencing of 84 diverse bacterial isolates was performed using an Oxford Nanopore Technologies PromethION 2 Solo using the Rapid Barcoding Kit 96 V14 (SQK-RBK114.96). Reads were basecalled with Dorado v0.9.5^10^ using the sup@v5.0.0 model and filtered to retain reads with mean quality *≥*10. Five isolates were selected, each from a different genus: *Enterobacter hormaechei, Klebsiella pneumoniae, Listeria innocua, Providencia rettgeri* and *Shigella flexneri*. Short-read Illumina sequencing was available for all samples and was used to polish the reference genomes (Table S1). Selection was based on high read depth and preliminary assessments showing no evidence of heterogeneity or divergence between the Illumina and ONT datasets.

For each genome, we followed our previously published method to generate a high-accuracy reference assembly. ^11^ Briefly, the ONT reads were assembled with Trycycler v0.5.5^9^ and the resulting genome was polished using Medaka v2.0.1^12^, Polypolish v0.6.0^13^ and Pypolca v0.3.1. ^14^ The resulting assemblies were highly accurate and used as ground truth.

ONT reads for each genome were divided into six non-overlapping 50*×* subsets for a total of 30 read sets. Each read set was assembled using the following long-read assemblers: Canu v2.3^15^, Flye v2.9.5^16^, LJA v0.2^17^, metaMDBG v1.1^18^, miniasm v0.3^19^, NECAT v0.0.1^20^, NextDenovo v2.5.2^21^, Raven v1.8.3^22^ and wtdbg2 v2.5^23^. Each of these tools was run via the helper scripts included with Autocycler, which included extra processing for Canu (overlap-trimming and repeat/bubble removal), metaMDBG (low-depth contig removal), miniasm (polishing with Minipolish ^24^) and NextDenovo (polishing with NextPolish ^25^).

In addition, we assembled each read set with the Dragonflye v1.2.1^26^ and Hybracter v0.11.2^27^ long-read assembly pipelines and the consensus assembly tool MAECI ^28^ (commit f1eb3d7). For Autocycler, we produced an automated assembly (using its autocycler full.sh script) and a manually curated assembly.

We also attempted to evaluate MAC2.0^29^, but it did not perform correctly on complete bacterial genomes, producing outputs with duplicated sequences. Other consensus assembly tools, such as quickmerge ^30^ and Metassembler ^8^, are older and were primarily designed to improve contiguity in fragmented eukaryotic assemblies. These tools are not suitable for refining complete bacterial genomes from long-read data and were therefore excluded from this comparison. ^31^

Each assembly was compared to its corresponding ground-truth reference using a custom script (`assess_assembly.py`) that aligns the assembly to the reference sequence with minimap2 v2.28^32^ and quantifies accuracy metrics including sequence errors (substitutions and indels), missing bases and extra bases. We also assessed assembly accuracy with Inspector v1.3.1^33^ and CRAQ v1.0.9^34^, which evaluate assemblies based on read alignments rather than a reference. Full commands are provided in the supplementary data.

### 3.2 Results

Among the single-tool assemblers, Canu and Flye consistently produced the fewest sequence-level errors, typically with fewer than 10 substitutions and indel errors per assembly (Figure 2A, Table S2). All other long-read assemblers had higher error rates. Across all tools, structural inaccuracies were common (Figure 2B), with assemblies often missing genomic elements (e.g. small plasmids) or containing spurious extra sequence (e.g. duplicated ends of circular contigs).

**Fig. 2.**
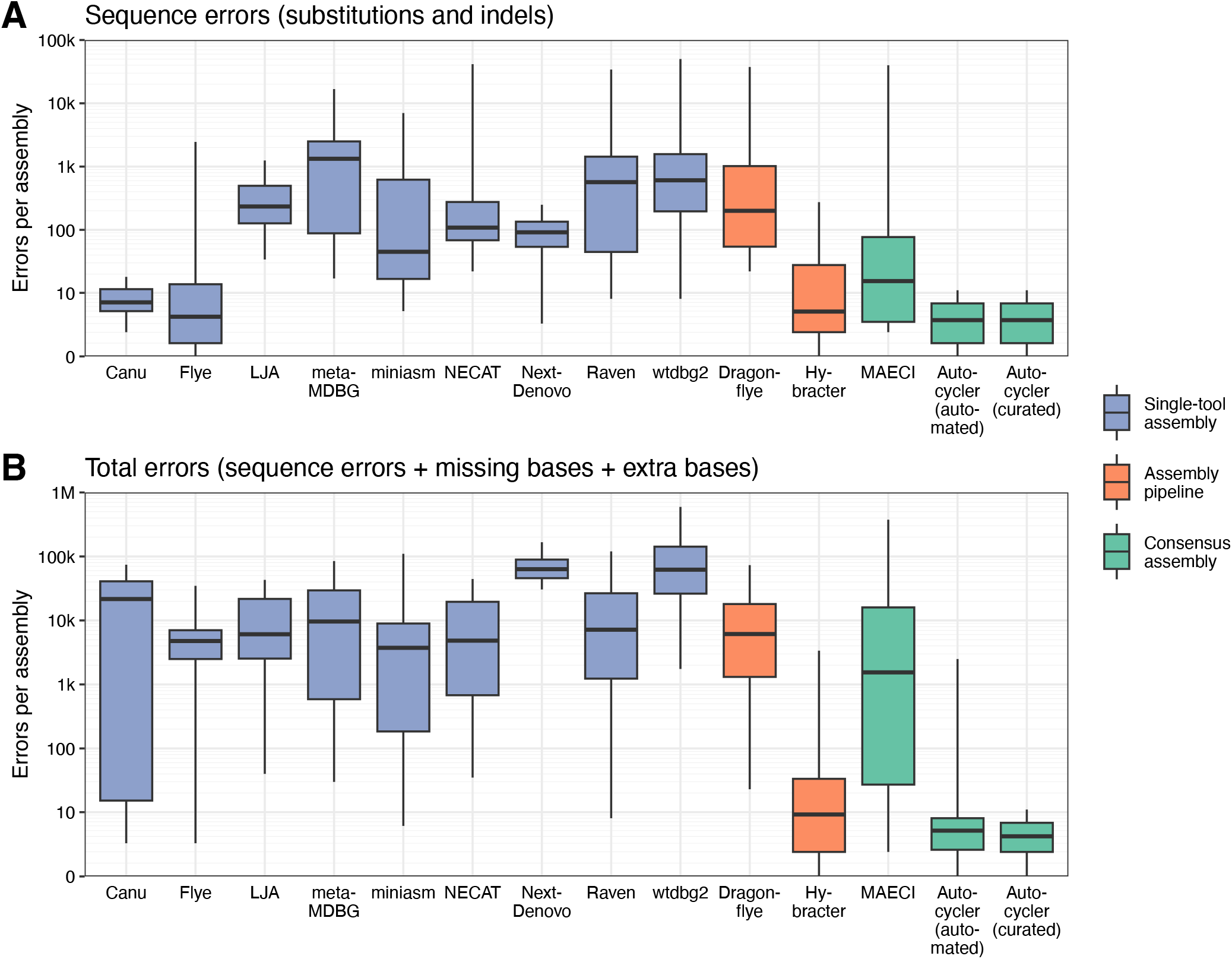
Assembler benchmarking results from the `assess_assembly.py` script. **(A)** Sequence errors (substitutions and indels); **(B)** Sequence errors and structural assembly errors (missing and extra bases). Lower values on the y-axes indicate better assembly accuracy. Results are coloured by category: individual long-read assembly tools (blue), long-read assembly pipelines (orange) and consensus assembly tools (green). Autocycler results are shown separately for fully automated and manually curated assemblies. Boxplot whiskers extend to the minimum and maximum values. The y-axes use a pseudo-logarithmic scale that accommodates zeros.

Of the long-read assembly pipelines, Hybracter outperformed Dragonflye. By integrating Plassembler ^35^, Hybracter improves plasmid recovery and avoids structural errors such as duplication of plasmids. However, for the *Enterobacter* and *Klebsiella* genomes, Hybracter showed elevated error rates in large plasmids, likely due to errors introduced during plasmid assembly by Unicycler ^36^ within Plassembler (Table S3). Dragonflye performed worse than Flye alone, likely due to its default use of Racon polishing (instead of Flye’s internal polisher) and the `--nano-raw` option which is suboptimal for modern ONT reads. With adjusted parameters (`--racon 0 --opts` ‘-i 1’`--nanohq`), Dragonflye?s performance matched that of Flye (Table S4).’MAECI was the only consensus assembly tool tested apart from Autocycler. Despite incorporating three input assemblers (Canu, Flye and wtdbg2), MAECI did not consistently outperform Canu or Flye.

When run in a fully automated manner, Autocycler produced assemblies with the lowest sequence error counts of any method (median: 3.5 errors per assembly, range: 0−11). It was also structurally accurate in most cases, successfully recovering all replicons for four of the five genomes. This is in part because `autocycler_full.sh` uses Plassembler when generating input assemblies. However, the smallest plasmid of the *Enterobacter* genome (2.5 kbp) was occasionally missed. In manually curated runs, the missing plasmid could be identified by inspecting the clustering tree (output by `autocycler cluster`) and included in the final assembly by overriding the default clustering. These curated Autocycler assemblies had no structural errors reported by `assess_assembly.py`, Inspector or CRAQ (Table S2).

## 4. Discussion and conclusions

Consensus assembly offers a clear accuracy advantage over single-tool assembly. In our benchmarking, even the best-performing individual assemblers (Canu and Flye) consistently made avoidable errors. Consensus approaches mitigate such issues by averaging over multiple inputs, reducing both small-scale errors and structural inaccuracies. Trycycler ^9^ provided a robust framework for generating consensus bacterial genome assemblies, but it requires substantial user intervention, limiting its scalability. Autocycler brings the benefits of consensus assembly into a fully automated workflow, enabling accurate bacterial genome assembly at scale.

Autocycler does not guarantee perfect results, as its consensus reflects the input assemblies. If the inputs are fragmented (e.g. due to a repeat longer than the read length), Autocycler cannot produce a fully resolved consensus. When most inputs share the same error, that error can persist into the final assembly. In our evaluation, there were typically fewer than 10 sequence errors (substitutions and indels) per Autocycler assembly (Table S5), most commonly homopolymer-length errors resulting from systematic basecalling issues in ONT reads. ^37^ The frequency of these errors depends on factors such as pore type (R10.4.1 is more accurate than R9.4.1), basecalling model (sup is more accurate than hac or fast) and bacterial strain. To address these errors, short-read polishing can be applied after Autocycler, yielding hybrid assemblies with maximal accuracy. ^14^

The only structural error observed in automated Autocycler assemblies in this study was the omission of a small plasmid, which many long-read assemblers fail to recover. ^6,38^ Autocycler supports manual intervention at key steps in its pipeline, enabling users to review the data and apply their own judgement to correct such issues. This flexible design allows Autocycler to function as both a scalable automated tool and as a framework for high-accuracy reference genome assembly.

Although not evaluated in this study, linear replicons pose additional challenges for genome assembly. Assemblers may erroneously extend hairpin ends or terminate open ends inconsistently. While Autocycler includes logic to detect and trim hairpin overlaps, full resolution of linear sequences still frequently requires manual intervention. Improved support for such cases remains an area for improvement for both long-read assemblers and Autocycler. Autocycler is open-source, well documented and easy to install. It requires only modest system resources (excluding input assembly generation) and provides intermediate outputs to support transparency and manual curation. It fills a key gap in the current assembly tool landscape: existing consensus tools either underperform or do not scale, while long-read assembly pipelines rely on a single assembler and inherit its limitations. Because it relies on multiple inputs, an Autocycler-based pipeline is more computationally intensive than using a single assembler, but it often yields better assemblies. We therefore recommend Autocycler for long-read bacterial genome projects where maximum assembly accuracy is required.

## 6. Author contributions statement

RRW conceived and programmed Autocycler with input from TPS. RRW performed the benchmarks and analysed the data. RRW, BPH and TPS wrote and reviewed the manuscript.

## 8. Data availability

Supplementary figures, tables and methods are available at github.com/rrwick/Autocycler-paper. Assemblies, reference genomes and read sets used in the analysis are available at figshare.unimelb.edu.au/projects/Autocycler/247142.

## 9. Acknowledgements

This research was performed in part at the Centre for Pathogen Genomics Innovation Hub, Department of Microbiology and Immunology, University of Melbourne at the Peter Doherty Institute for Infection and Immunity.

This paper acknowledges the PulseNet Asia-Pacific team at the Centre for Pathogen Genomics and the Microbiological Diagnostic Unit Public Health Laboratory (MDU PHL) for contributing data and isolates to the study. Support for PulseNet Asia-Pacific is funded by the US Centers for Disease Control and Prevention (CDC) Global Antimicrobial Resistance Laboratory and Response Network through the Association of Public Health Laboratories. MDU PHL is funded by the Victorian Government, Australia.

We thank the Autocycler alpha testers for their valuable feedback prior to the tool’s public release: Alex Krause, Bogdan Iorga, Dan Whiley, Danielle Ingle, Erin Young, George Bouras, Josh Zhang, Mariel Beiers, Marko Verce, Matthew Croxen, Mona Taouk, Munazzah Maqbool, Oliver Schwengers, Sarah Baines, Steve Baeyen, Sudaraka Mallawaarachchi, Tatum Mortimer, Tue Sparholt Jørgensen, Tung Trinh and Ying Xu.

