## Supplementary figures and images for "Autocycler: long-read consensus assembly for bacterial genomes"

### Figure S1

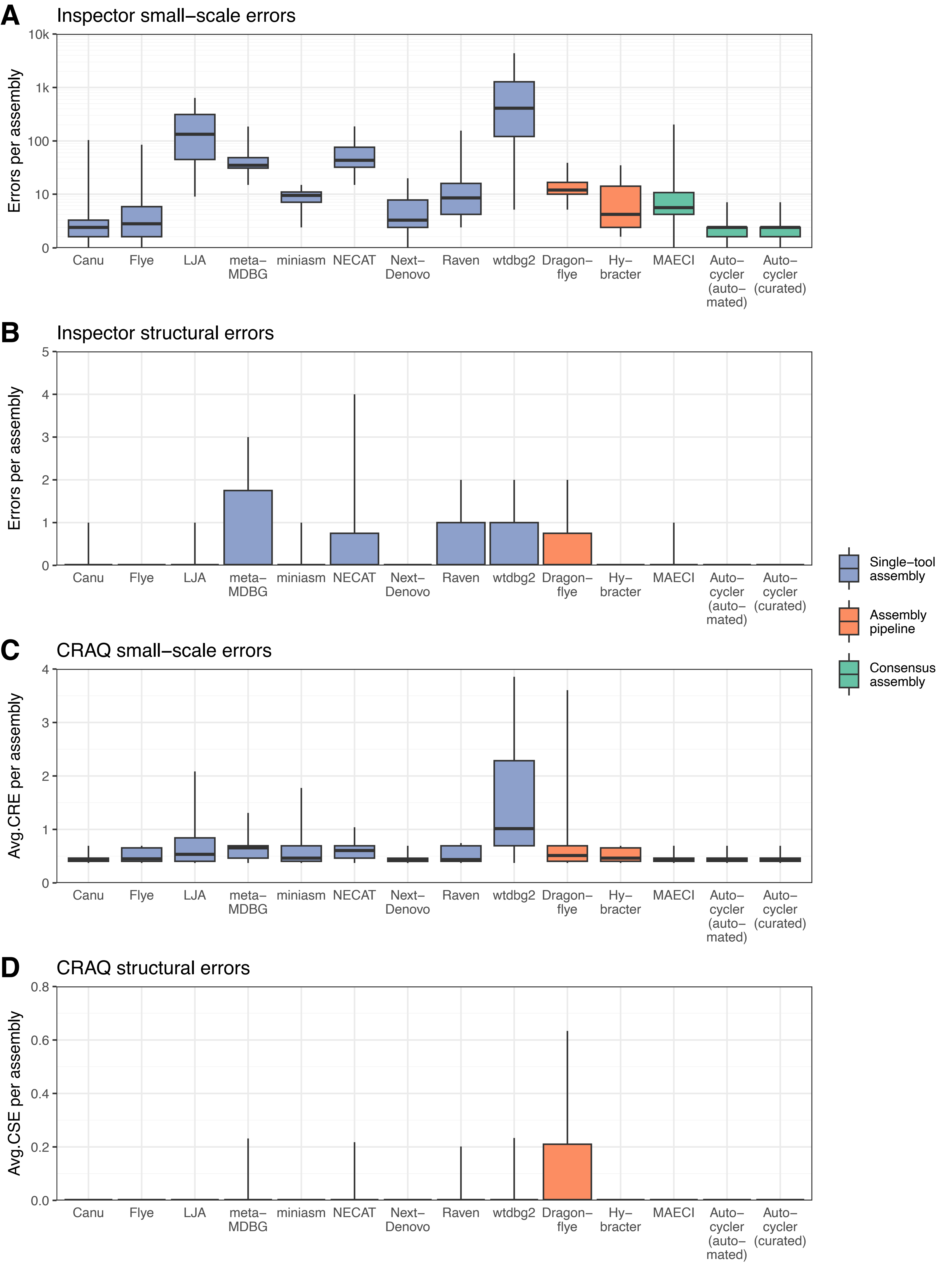
